# Voltage gated N-type Ca-channels in Neurogliaform interneurons in the rat Prefrontal Cortex

**DOI:** 10.1101/299503

**Authors:** Nils Ole Dalby, Morten Grunnet

## Abstract

The fastspiking parvalbumin expressing basketcells (FS BC) and the neurogliaform interneurons (NGF IN) innervate somatic and dendritic compartments of postsynaptic pyramidal neurons, respectively. Here we have used paired recordings of each type of interneuron with a postsynaptic pyramidal neuron in layer II-IV of the adult rat prefrontal cortex (PFC) to characterize the interneuron action potentials (AP), evoked IPSC characteristics and response to the GAT-1 inhibitor NNC 711 and the N-type Ca-channel inhibitor ω-conotoxin GVIA. GABA released from FS BCs were insensitive to ω-conotoxin GVIA (0.5 μM), the postsynaptic response did not desensitize and the IPSC was unchanged in NNC 711 (2 μM). Conversely, IPSCs generated by APs from NGF INs were 80 % inhibited by ω-conotoxin GVIA, displayed a strong degree of desensitization and the decay-time of the IPSCs doubled in NNC 711. We used these pharmacological differences to identify the source of GABA in paradigms of synaptic overspill induced by extracellular stimulation. The decaytime of the electrically evoked event doubled in NNC 711 and this increase was reversed by subsequent application of ω-conotoxin GVIA, suggesting that the source of GABA responding to NNC 711 originate from axons that terminate on pyramidal neuron dendrites and employ N-type Ca-channels for release.

## Introduction

Cortical GABAergic interneurons can be categorized into three major groups based on the expression of either the Ca-binding protein parvalbumin (PV), the neuropeptide somatostatin (SST) or the 5HT_3a_-receptor (reviewed by (Rudy B et al. 2011)). These groups can be further subdivided based upon (among others) action potential (AP) properties, frequency of APs, dendritic or somatic innervation and characteristics of evoked IPSCs in the target neuron. Two types of interneurons in the prefrontal cortex (PFC) with a different innervation pattern onto postsynaptic pyramidal neurons are the PV-expressing fast spiking basket cells (FS BC), which innervate the perisomatic regions to deliver fast activation of predominately α_1_-containing synaptic GABA_A_ receptors, and the more recently characterized 5HT_3A_-R expressing neurogliaform interneurons (NGF IN), which exert a slow inhibition at dendritic loci, mediated by both synaptic and extrasynaptic GABA_A_ and GABA_B_ receptors (Tamas G et al. 2003; Freund TF and I Katona 2007; Olah S et al. 2007; Szabadics J et al. 2007; Olah S et al. 2009). The PV-containing FS BCs transmit in a manner characterized as point-to-point transmission onto pyramidal neurons whereas the release from NGF INs is more aptly described as volume transmission, based on the large distance between release-site and postsynaptic receptor (~2.7 μm, (Olah S *et al.* 2009)) which, together with interneuron firing frequency, also predict different roles for the interneurons in network function (Fuentealba P et al. 2008; Sohal VS et al. 2009; Karayannis T et al. 2010; Chen G et al. 2017). Volume transmission of GABA from the NGF INs reach extrasynaptic δ-subunit containing GABA_A_ receptors, a population of receptors which also can be activated by evoking synaptic overspill. By evoked synaptic overspill is usually understood the concerted release evoked by a local extracellular stimulation electrode from terminals in vicinity of the field, leading to overspill of GABA from the synapse in amounts that surpass the buffering capacity of the GABA transporter-1 (GAT-1) to activate extrasynaptic receptors (Overstreet LS and GL Westbrook 2003). The evoked overspill to extrasynaptic receptors is greatly enhanced by inhibition of GAT-1, and this characteristic is shared with GABA release from NGF INs onto pyramidal neurons (Szabadics J *et al*. 2007). This also illustrate that the effect of inhibitors of GAT-1 in brain slices is measured indirectly, by the following activation of extrasynaptic GABA_A_ and GABA_B_ receptors. For this reason, attempts to delineate the individual components in response to changes in the response to GAT-1 inhibition in, for example, paradigms of cognitive impairment, points to several possible reasons (Gonzalez-Burgos G et al. 2009; Kjaerby C et al. 2014; Frankle WG et al. 2015). Therefore, we have in the present study aimed at characterizing in more detail the role of GAT-1 in GABAergic transmission in the layer II-IV of the adult rat prefrontal cortex (PFC). Using paired recordings of FS BC – pyramidal neurons and NGF IN-pyramidal neurons, we demonstrate that inhibition of GAT-1 enhance the decaytime of the NGF IN IPSC only, with no effect on FS-BC IPSCs. Further, we demonstrate that GABA release from the NGF INs is effectively blocked by the N-type Ca-channel blocker ω-conotoxin GVIA, which is ineffective at FS BC-pyramidal neuron synapses, known to employ the P/Q type Ca-channels. Finally, we combine these experiments to demonstrate that synaptic overspill, induced by electrical stimulation and inhibition of GAT-1, is reversed by ω-conotoxin GVIA at similar concentrations that block release from NGF INs, thus strongly supporting that GAT-1 mediated effects are not somatic but dendritic in origin, and due to release from interneuron terminals employing N-type Ca-channels for GABA release.

## Methods and Materials

### Animals

Young adult male Sprague Dawley rats were obtained from Envigo (Holland) and weighing 250-300 g at the time of experiment. Animals were housed in cages holding up to 6 rats at a normal 12 h/12 h light/dark cyclus with ad libitum access to food and water. Animal experiments were carried out in accordance with the European Communities Council Directive (86/609/EEC) for the care and use of laboratory animals and the Danish legislation regulating animal experiments.

### Slice preparation

Rats were decapitated using a guillotine and the head immediately immersed in icecold N-methyl-d-glucamine (NMDG)-kynurenate-artificial cerebrospinal fluid (ACSF) of the composition (in mM): N-Methyl-D-Glucamine (NMDG)-HCl (100), NaCl (26), NaHCO_3_ (26), Glucose (11), KCl (2.5), NaH_2_PO_4_ (1.25), CaCl_2_ (1), MgCl_2_ (3), pyruvate (1), ascorbate (0.3) and kynurenic acid (2). All ACSF solutions was adjusted to osmolarity of 310 +/-3, was aerated with carbogen and held a pH of ~ 7.4. The head was cooled for ~ 5 minutes before dissection of the brain began. The frontal cortex (rostral part of brain after coronal cut from approximately bregma) was isolated and glued to the platform on a Leica VT1200 S vibrating microtome and cut in 350 μm thick coronal sections in icecold NMDG-kynurenate-ACSF. Slices were stored for 1-5 h at 28 - 29 °C in ACSF of similar composition as the cutting solution, but with NMDG replaced by equimolar NaCl and omission of kynurenic acid. Slices were used for recordings no earlier than 1 h after cutting. Slices were not hemisected and the PFC was contained in typically 4 slices.

### Recordings

Slices were transferred to the recording chamber and perfused at a flowrate of 2.6 – 2.8 ml/min at 33-34 °C with ACSF of a similar composition as storage-ACSF but with following changes: CaCl_2_ (2), MgCl_2_ (2), kynurenic acid (2), CGP54626 (0.001), LY341495 (0.001). Suitable pairs of pyramidal neuron – interneuron localized in the layer II-IV of the PFC were visualized under IR illumination. Recording pipettes were made of borosilicate glass (OD/ID 1.5/1.1 mm, Sutter) with a resistance of 8-9 MOhm for interneurons and 6-8 MOhm for pyramidal neurons. Interneurons were chosen based on morphology and patched with a KMeSO_3_ based intracellular solution of the composition (mM): KMeSO_3_ (135), KCl (4), NaCl (4), MgCl_2_ (2), EGTA (0.2), HEPES (10), ATP (2), GTP (0.5), pH 7.2 to determine their subtype, assessed from firing frequency and AP waveform. A suitable pyramidal neuron located within 10 – 30 μm from the interneuron was patched next using a CsCl based intracellular solution of the composition CsCl (135), NaCl (4), MgCl_2_ (2), EGTA (0.1), HEPES (10), ATP (2), GTP (0.5), QX-314 (5), TEA (5), pH 7.2. Osmolarity of intracellular solutions was adjusted to 292 mOsm. Recordings began no earlier than 6 minutes after successful patch of the postsynaptic neuron. Interneurons held in current clamp were compensated for electrode capacitance and bridge balanced and pyramidal neurons in voltage clamp were 70 % compensated for series resistance (RS). Neurons that changed more than 30 % in RS (postsynaptic) or bridge balanced resistance (presynaptic) during recordings were excluded from analysis. Recordings were made using a Multiclamp 700A patch clamp amplifier controlled by pClamp 9 software and digitized at 20 kHz with a digidata 1322 (all Molecular Devices) and filtered (4-pole Bessell) at 5 kHz. First a series of rectangular current input (-87 to 500 pA for 400 ms in steps of 25 pA) was delivered to the presynaptic interneuron at a sweep interval of 12 s. For pharmacological analyses of GAT-1 and ω-conotoxin GVIA we used single evoked APs by 800-1000 pA input current for 2.5 ms at 30 s sweep interval. Stimulation electrodes used for single evoked IPSCs was pulled from ϴ-glass (OD 1.5 mm, Sutter), to an opening of 3-5 μm and filled with ACSF. The evoked events commenced at the earliest of 6 minutes after establishing whole-cell, and baseline recording began after 3 minutes of stable responses.

### Analysis

The passive characteristics of the presynaptic interneurons measured were membrane potential, input resistance and membrane time constant measured as the mean of mono-exponential fits to the first two hyperpolarizing steps. The action potential (AP) characteristics were determined as averages of all APs in a sweep. Parameters were AP peak, after hyperpolarization (AHP) peak, rheobase (current at first AP threshold voltage) and halfwidth of AP (FWHM: full width at half-maximal amplitude). All APs were isolated in pClamp using the event threshold function, sampling 1 ms before and 4 ms after the peak. Traces were analyzed in OriginPro 2016 using routines written in Origins LabTalk programming language. The threshold voltage is determined as a local maximum of d^3^V/dt^3^ (Figure 1C, (Henze DA and G Buzsaki 2001)). All values of AP parameters were analyzed on the interpolated traces, points connected by b-spline. The inhibitory postsynaptic current (IPSC) was analyzed as total charge of the postsynaptic trace during on- and offset of the presynaptic input current. For the IPSCs evoked by single stimulation of the presynaptic IN, or by stimulation electrode, the decay was analyzed as the weighed decay, τ_W_ (Charge_peak_/peak) to obviate cell-to-cell variability in number of exponentials that best fit the decaytime. A comparison of this measure to a single exponential fit in baseline in the case of paired recordings, is given in table 2. All data throughout are presented as the mean ± SD or as range where indicated. The spontaneous IPSCs were analyzed in minianalysis 6.03 (Synaptosoft) as previously detailed ((Kjaerby C *et al.* 2014).

**Figure 1.**
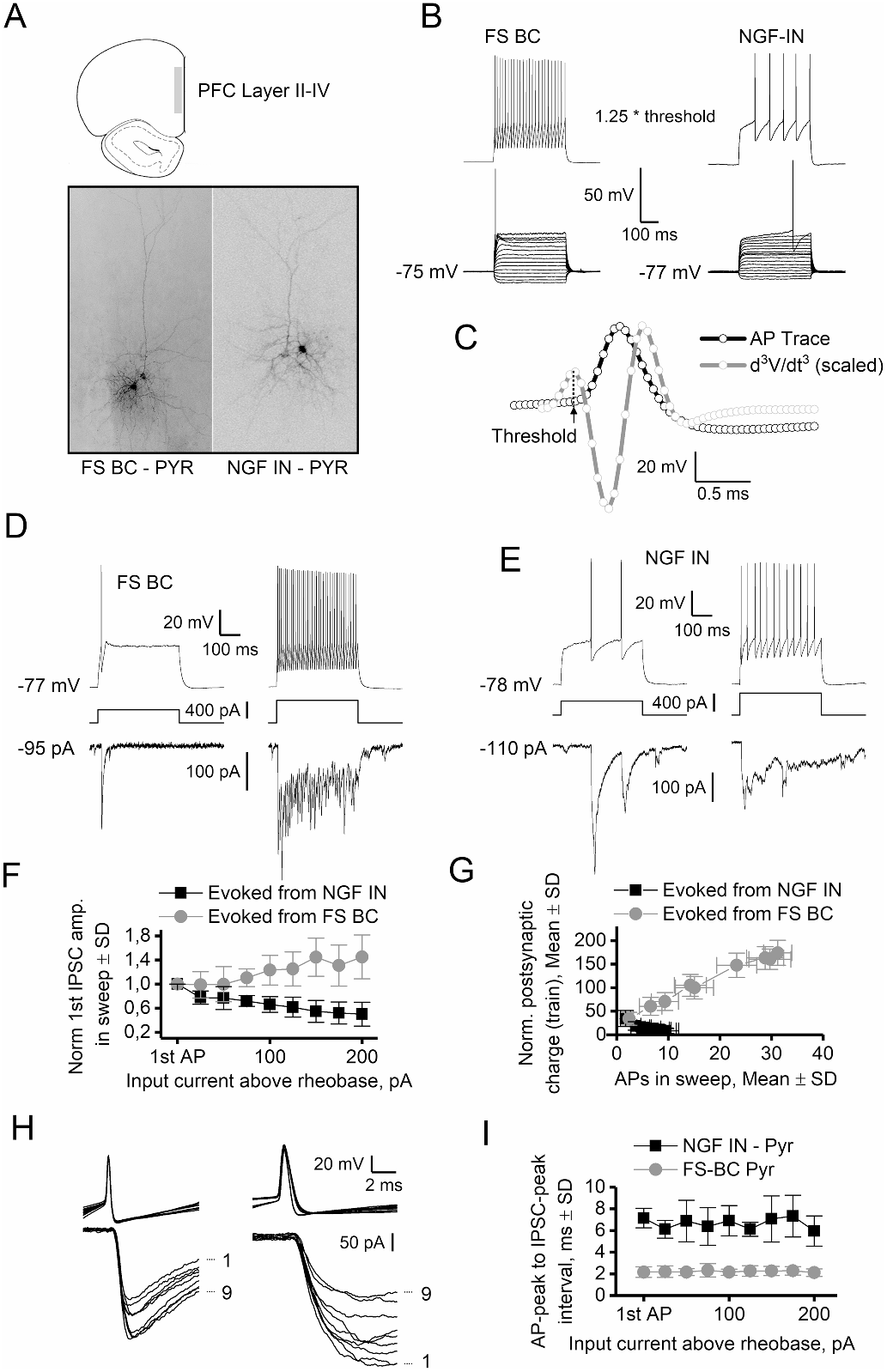
FS-BC and NGF INs in the layer II-IV of the adult rat PFC. **A**: Pairs of FS BC-pyramidal neuron (left) and NGF IN-pyramidal neuron (right) in layer II-IV. **B**: Initially, passive and active properties of interneurons were characterized in a rectangular step-protocol (400 ms duration, 25 pA step). FS-BC and NGF-INs both had similar low membrane time constant (6-7 ms), V_m_ < -75 mV and 110 - 130 MΩ input resistance, but differed very significantly in the AP properties, frequency and IPSC characteristics evoked in the postsynaptic pyramidal neuron. AP properties shown in table 1 were determined as the average of the APs in the sweep closest to 1.25^∗^ above the threshold current for the first AP. **C**: The threshold voltage (red horizontal line) is determined as a local maximum of the 3^rd^ derivative of the AP (red vertical line). The d^3^V/dt^3^ trace is scaled in amplitude for the purpose of illustration (scalebar is for the AP, black trace). Digitization points of the trace are marked as small circles and FWHM is indicated as a red line in the peak. **D-E**: Single AP and train APs in pair of FS BC-pyramidal neuron (**D**) and NGF IN-pyramidal neuron pair (**E**). **F-G**: A strong use-dependent desensitization of the NGF IN-pyramidal neuron synapse(s) was observed. In a step protocol with a sweep interval of 12 s, the 1^st^ IPSC and the train IPSC response, despite an increase in number of APs with input current, would both diminish in amplitude and charge with frequency of use. The FS-BC pyramidal neuron synapse displayed instead an increase in the 1^st^ AP amplitude and charge of the evoked IPSC train with increasing input current and frequency of presynaptic APs. The charge of the train of IPSCs (**G**) was normalized to the peak of the first IPSC in the train. n = 12 NGF IN-Pyr pairs and 14 FS BC-Pyr pairs. **H-I**: IPSC rise time and AP-peak to IPSC-peak is unchanged for the first IPSC in the train from 9 sweeps of increasing current input despite significant change of amplitude. Numbers 1 and 9 (marked next to IPSCs in I) refer to the first sweep with an AP and the 8 following sweeps at increasing input current and synaptic use (numbers of APs generated). Same pairs as figure **F-G**.

**Table 1.** Passive and active properties of FS-BC and NGF INs subjected to rectangular current steps.

|  | FS BC (n=14) | NGF IN (n=12) |
| --- | --- | --- |
|  | Mean ± SD |  |
| V<sub>m</sub>, mV | -75.3 ± 3.1 | -76.0 ± 3.3 |
| Resistance, MΩ | 111.9 ± 35.5 | 125.3 ± 28.8 |
| Membrane time constant, τ<sub>m</sub> | 6.4 ± 1.7 | 6.7 ± 1.3 |
| AP threshold, mV | -41.7 ± 4.2 | -32.9 ± 5.6 |
| Rheobase, pA | 338 ± 76 | 276 ± 49 |
| AP Peak, mV | 58.5 ± 6.4 | 54.0 ± 4.9 |
| FWHM, ms | 0.35 ± 0.06 | 0.70 ± 0.10 |
| AHP, mV | 20.9 ± 1.8 | 22.1 ± 3.1 |

## Results

### Initial characterization of FS BC and NGF INs in the adult rat PFC

The passive and active properties of FS BC, NGF IN and the evoked inhibitory postsynaptic train in synaptically connected pyramidal neurons was determined from 14 and 12 pairs, respectively and presented in figure 1 and table 1. These recordings were not used for other analyses presented later. All pairs of IN-pyramidal neurons were located in the rat PFC layer II-IV and had distances between somas of 10 – 50 μm (range), and at similar distance to the pial surface (paired neurons were from the same cortical layer). A few pairs were filled with biocytin and stained with streptavidin coupled Alexa-488 (EGFP) to illustrate localization and neuronal arborization (figure 1A). FS-BC and NGF INs were patched in voltage clamp (VC) mode and all displayed a positive holding current upon entry (V_h_ -70 mV) and had capacitances of 8 – 10 pF (range). In current clamp (CC) mode the resting V_m_ (mean±SD) was -75.3±3.1 mV (FS-BC) and -76.0±3.3 (NGF IN), indicating the presence of a considerable leak, presumably 2-pore domain K-channels. Following establishment of a stable whole cell configuration of the IN in CC, a pyramidal neuron near was patched. The success rate for patch of a connected postsynaptic pyramidal neuron appeared to increase with the depth of particularly the pyramidal neuron, which was typically located 10-20 μm deeper than the IN. Pyramidal neurons had capacitance of 11-15 pF and RS of 15–25 MΩ (range). FS BCs and NGF INs were similar in passive properties, but differed very significantly in AP properties and postsynaptic response (see table 1 and figure 1). The rheobase for the AP in FS BCs was (mean±SD) 338±76 pA and most cells displayed a stuttering firing behavior just above this threshold. A non-accommodating firing pattern produced (rounded mean±SD) 12±9 and 40±19 APs in the 400 ms depolarization window at input currents of 1.25 and 1.75^∗^rheobase, respectively (figure 1B, left). The NGF INs always displayed a late generation of the first AP after a slow rise of the depolarization plateau to reach threshold voltage (figure 1B, right). The average rheobase for the AP in NGF INs was (mean pA±SD) 276±49 and a rounded mean±SD of 5±2 and 9±2 APs was produced at input currents of 1.25 and 1.75^∗^rheobase, respectively. We did not go higher than 2.5 ^∗^ rheobase input current, but never saw AP train frequencies > 35 Hz in NGF INs. At high input current, the AP amplitude was constant but frequency pattern was more accommodating due to absence of late-spiking (LS) phenotype of the first two-three APs. This characteristic is reminiscent of the LS1-type, described by (Miyoshi G et al. 2010). The evoked IPSCs in the postsynaptic pyramidal neuron had very different kinetics dependent on type of presynaptic IN, fast rise and decay for FS BC and slow rise and decay for NGF INs, characterized below. Interestingly, the train of IPSCs generated in the course of increased step-current (12 s between sweep), behaved differently in the time course of the experiment for the two types of pairs. For the FS BC–pyramidal neuron pair, we observed a steady increase in the amplitude of the first IPSC in the train with increasing input current (figure 1F) and also a near-linear increase in the charge of the entire IPSC train with increasing input current and number of APs in sweeps (figure 1G). For the NGF IN-pyramidal neuron pair, an opposite effect was observed. The amplitude of the first IPSC and especially charge of the train of IPSCs decreased fast with increasing number of APs in the sweeps (figure 1G, H), leading in most cases to an unresolvable postsynaptic response after the first IPSC at 2 – 2.5^∗^rheobase input current. This plasticity was partially reversed after a period of 8-10 min without presynaptic activity, but the IPSC charge was quickly lost again when presynaptic activity resumed (not shown). The extrasynaptic GABA_A_ receptors are a component of the IPSCs evoked from NGF INs, so we tested if the response to the orthosteric agonist THIP (2 μM) at δ-subunit containing GABA_A_ receptors in layer II-IV pyramidal neurons displayed signs of desensitization. At the end of a 6 min long exposure, the tonic current density (pA/pF±SD) in THIP was 3.5± 1.3 (n=7), and we did not observe a decrease in this response during wash-in, indicating that postsynaptic desensitization to an orthosteric agonist at δ-subunit containing GABA_A_ receptors did not occur. We also did not observe a reduction in the IPSCs amplitude in single evoked APs by a 2.5-ms step to 800-1000 pA at a sweep interval of 30 s, the protocol used to assess pharmacological effects (below). For these reasons, the rapid reduction in the charge of IPSC trains evoked from NGF INs appear use-dependent and presynaptic in origin. The reason for the increased response in the FS BC-pyramidal neuron synapse is less clear but may involve different release probabilities of individual release sites contributing to the composite IPSC. The IPSC rise time and time-to-peak was not affected by the change in IPSC peak for either type of pairs induced in the protocol (figure 1H, I).

### Single evoked IPSCs and effect of GAT-1 inhibition

Because of the observed plasticity of the NGF IN-pyramidal neuron synapse, we used single evoked APs with a 30 s long sweep interval for investigation of pharmacological effects. At this frequency, responses were stable for both types of pairs. Statistical assessment of drug-effect was made on IPSC-means of sweep 8-10 (baseline) and sweep 23-25 (effect). The baseline characteristics of single IPSCs evoked from 14 FS BC-pyramidal neuron pairs and 16 NGF IN-pyramidal neuron pairs is given in table 2. From these pairs, 7 FS BC-pyramidal neuron pairs and 8 NGF IN-pyramidal neuron pairs were used for assessment of the effect of GAT-1 inhibition (figure 2), and 7 FS BC-pyramidal neuron pairs and 8 NGF IN-pyramidal neuron pairs were used for assessment of the effect of ω-conotoxin GVIA (figure 4). The IPSC evoked from the FS BC had a baseline τ_W_ of 9.3±2.5 and 9.1±2.5 (mean ms±SD) in the presence of the GAT-1 inhibitor NNC 711 at sweep 23-25, the difference being ns (P=0.49, paired t-test, n=7). However, the τ_W_ of the IPSC evoked from NGF IN was 35.6±12.9 ms in baseline and 72.4±19.0 ms in 2 μM NNC 711 (P=3E-4, paired t-test, n=8). The averaged spontaneous event in the same neurons displayed a borderline significant increase (average 6 %) in the decaytime in NNC 711 (P=0.05, paired t-test, n=15, figure 2 C, D). The events evoked from the FS BC were similar in rise- and decay-kinetics to the spontaneous events, whereas the events evoked from the NGF INs had much longer rise- and decay-kinetics (figure 1 I, and figure 3 A, B, table 2). The *E*_Cl_ for the presynaptic IN was -64 mV, and in a few cases we saw single spontaneous slow depolarizing IPSPs in the interneuron (V_rest_ ~ -76 mV) and a simultaneous slow IPSC in the pyramidal neuron, not associated with the stimulation, thus most likely originating from neighboring NGF INs releasing GABA onto both neurons. However, stimulation of the presynaptic NGF IN was never associated with positive IPSPs in the interneuron i.e., we saw no examples of autapses. Several parameters could contribute to the different IPSC waveforms from FS BC and NGF INs, such as subtype of postsynaptic receptor and electrotonic distance between activated receptors and soma. We did not perform a detailed histologic analysis of the interneuron axonal termination on the pyramidal neurons, but we determined the IPSC reversal potential in 5 pairs of either subtype, not used for other pharmacological analyses (figure 2 E, F). The IPSC reversal potential, V_rev_, was (mean±SD) 2.1±5.6 mV for the FS BC-pyramidal neuron pairs and -14.9±4.7 mV for the NGF IN-pyramidal neuron pairs (P=0.001, t-test), strongly supporting a difference in the electrotonic distance to soma of receptors activated by GABA released from FS BC or NGF INs, most likely due to inadequate space clamp properties in the postsynaptic neuron.

**Figure 2.**
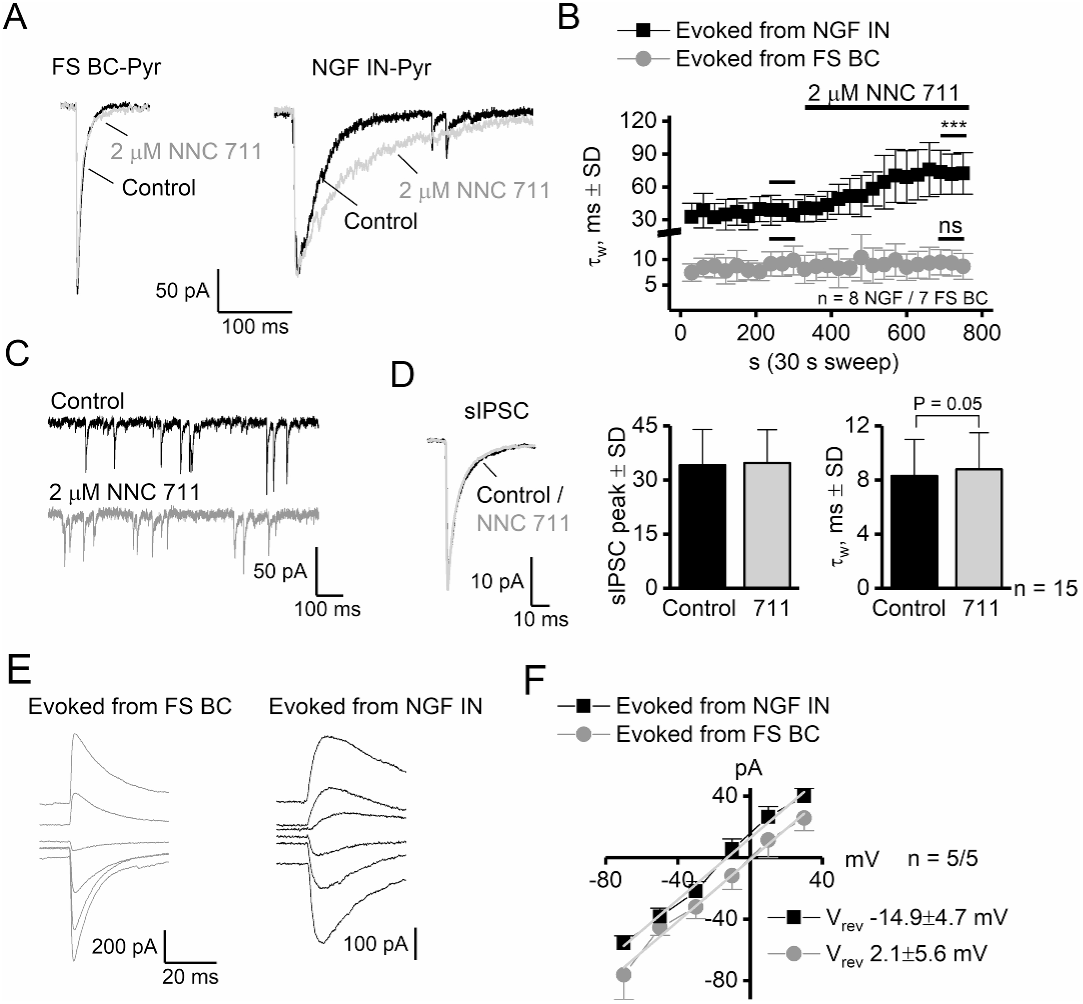
The FS-BC pyramidal neuron synapse and the NGF interneuron-pyramidal neuron synapse responded very differently to inhibition of GAT-1. **A**: Individually evoked IPSCs in pyramidal neurons show no effect of the GAT-1 inhibitor NNC-711 on the event evoked from the FS-BC but induce a doubling on the weight of tau, τ_W_ measured on IPSCs evoked from the NGF interneuron. **B**: Summary on the effect of NNC-711 on the IPSC decay time, determined from 8 NGF-pyr and 7 FS BC-pyr pairs. **C-D**: Representative traces of spontaneous activity from a pyramidal neuron in control and during exposure to 2 μM NNC-711. The averaged sIPSC waveform displayed a very small but borderline significant increase in the decaytime (paired t-test, P=0.05, n=15). **E-F**: The IPSC evoked from NGF INs reversed at a significantly more negative potential (-14.9±4.7 mV) than the IPSC evoked from the FS BCs (2.1±5.6 mV). The IV fit of the individual IPSCs was normalized to the conductance for the purpose of illustration.

**Figure 3.**
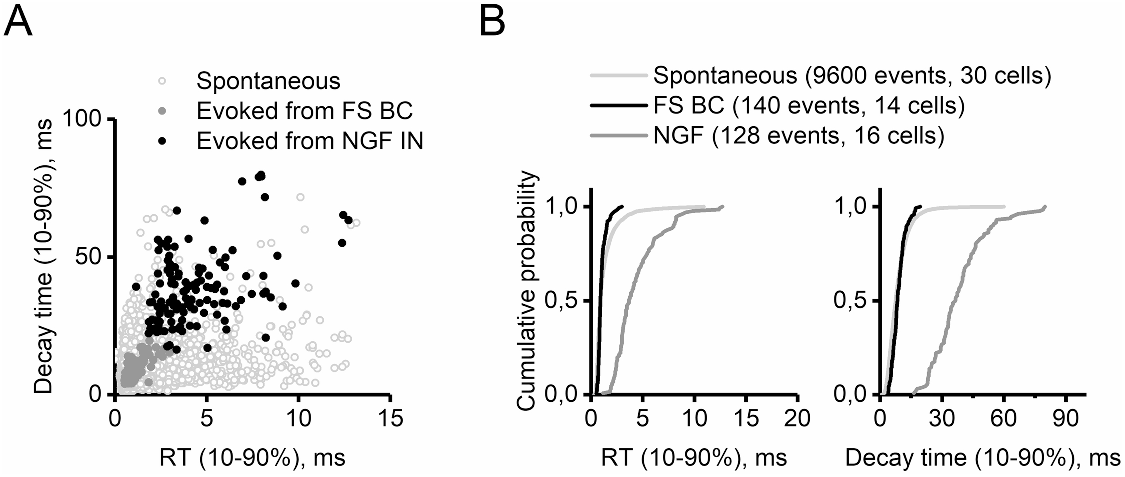
Kinetics of the evoked and spontaneous IPSCs. **A**: Scatter plot of the rise- and decay times of sIPSCs (9600 events, 320 events from each of 30 pyramidal neurons), 140 events evoked from FS-BC (14 pairs, 10 events each) and 128 events evoked from NGF interneurons (16 pairs, 8 events each). All events were from baseline recording period from tests of NNC-711 or ω-conotoxin GVIA. **B**: Probability density plots for the same events.

**Table 2.** IPSC characteristics of single evoked IPSCs from FS BC and NGF IN at 30 s interval

|  | Evoked from FS BC (n=14) | Evoked from NGF IN (n=16) |
| --- | --- | --- |
| | Mean ± SD (range) | |
| Amplitude, pA | 327 ± 163 (54 - 602) | 215 ± 217 (66 - 640) |
| RT (10-90%), ms | 0.9 ± 0.2 | 4.5 ± 2.0 |
| Decay time (single exp fit), ms | 7.7 ± 1.9 | 31.5 ± 15.0 |
| Weight of tau, $\tau_w$, (charge<sub>peak</sub>/peak), ms | 9.3 ± 2.5 | 35.6 ± 12.9 |

### Effect of N-type (Ca_v_2.2) Ca-channel blocker ω-conotoxin GVIA

The development of different types of postsynaptic response to trains of APs evoked from NGF IN and FS BCs, in a manner that did not alter the IPSC kinetics (figure 1 I, J), could indicate different functionality or type of presynaptic voltage gated Ca-channels in the FS BC and the NGF INs. The pore of N-type (Ca_v_2.2) voltage gated Ca-channels is selectively and potently blocked by ω-conotoxin GVIA, venom from the cone snail *Conus geographus* which is used to pharmacologically distinguish P/Q type (Ca_v_2.1) from N-type Ca-channels. Perfusion of 0.5 μM ω-conotoxin GVIA led to a rapid diminishment of the IPSC evoked from NGF INs, but had no effect on the IPSC evoked from FS BCs (figure 4 A-D). Assessment of the drug effect made on the average of sweep 8-10 (baseline) against average of sweep 23-25 (effect) for the NGF IN-pyramidal neuron synapse gave P=0.001 (paired t-test, n=8), an average of 80 % reduction of the evoked IPSC amplitude (range 71 - 96 % reduction). The effect of ω-conotoxin GVIA on the FS BC-pyramidal neuron synapse was ns (P=0.43, paired t-test, n=7). Scaling of two IPSCs per recording that were reduced in amplitude by at least a factor 3 (range 3.5 - 5.3, n = 8 pairs) by ω-conotoxin GVIA compared with two control IPSCs, indicated that the waveform was unaffected by presynaptic N-type Ca-channel blockage (the mean τ_W_ ± SD in control IPSCs was 30.1±14.7 ms and 29.7 ±13.9 ms in the peak scaled IPSCs recorded in ω-conotoxin GVIA, P=0.87, paired t-test, n=8, figure 4 C, D). This suggested that the NGF IN-pyramidal neuron IPSC kinetics was not due to asynchronous release from the NGF IN terminals. In addition, although the IPSCs evoked from NGF INs in ω-conotoxin GVIA were not mini-IPSCs given that failures were not reliably observed, it may be argued that detected spontaneous release from a NGF IN, would be similar in kinetics to the IPSC evoked from the same NGF IN. We saw no effect of ω-conotoxin GVIA on the peak of the averaged spontaneous event (paired t-test, P=0.16, n=7+8, combined for presynaptic NGF IN and FS BC pyramidal neuron pairs, figure 4 F).

**Figure 4.**
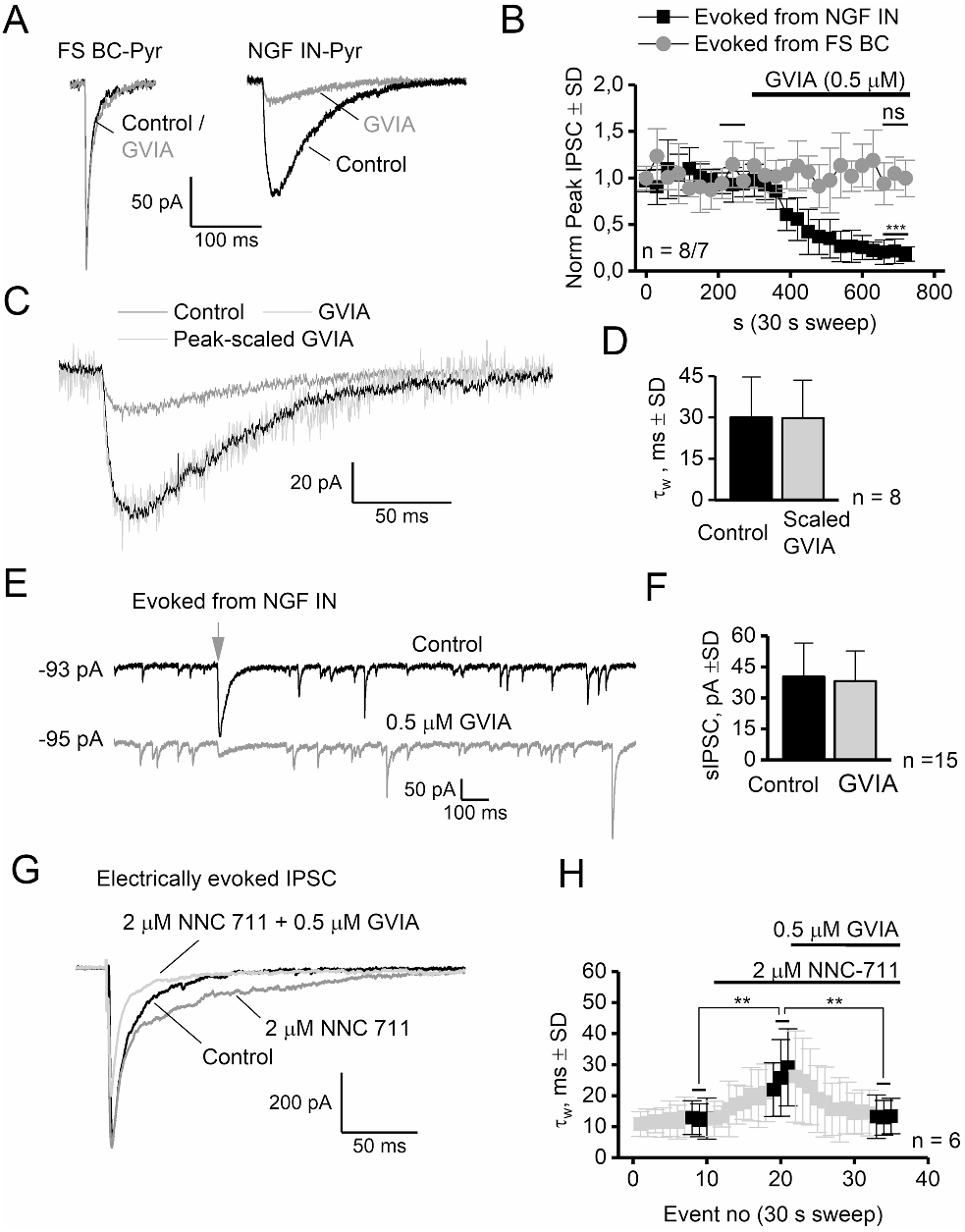
The N-type Ca-channel inhibitor ω-conotoxin GVIA (0.5 μM, GVIA in short notation) significantly reduce the amplitude of IPSCs evoked from NGF INs, but has no effect on IPSCs evoked from FS BCs. **A**: Traces of IPSCs evoked from FS BCs or NGF IN in control conditions or GVIA. **B**: Summary of the IPSC peak values in pyramidal neurons evoked from either type of IN. GVIA reduce the amplitude of IPSCs evoked from NGF INs with 80 %, assessed on mean response from sweep 8-10 (BL) and sweep 23-25 (GVIA), ^∗∗∗^: P=0.001, paired t-test, n=8. **C-D**: Events that was 70 % - 85 % reduced by GVIA (scale factor 3.5 – 5.2, range, n=8), was peak-scaled and shown to correlate well with the control IPSC. Black highamplitude trace is the control IPSC, grey low-amplitude trace is the event evoked in GVIA and the highamplitude noisy trace is the GVIA trace peak-scaled by the ratio of the control and GVIA trace. **E-F**: Representative traces showing spontaneous IPSCs and the IPSC evoked from NGF INs in control and presence of GVIA. The averaged spontaneous event was not reduced in GVIA. **G-H**: IPSCs electrically evoked with a bipolar stimulation electrode 40-50 μm from the soma of a pyramidal neuron display a significant increase in decay time in NNC 711, an effect which was completely removed in GVIA. Assessment of drug-effects was made on three events in each period (black squares) using one-way ANOVA/Bonferroni (^∗∗^: P<0.004, n=6, see text for details).

Because of the similarity in response to both NNC 711 and GVIA of the IPSCs evoked from FS BCs and the spontaneous events, we suspected a sparse representation of non-evoked NGF INs events in the population of spontaneous events. We performed a discriminant analysis using the paired events rise- and τ-values as variables to train a model to predict the origin of the spontaneous events (OriginPro 2017 software). This analysis was made on the Log_10_ values of RT and τ-values to transform the skewed distribution into a Log(normal) distribution for which normality was not rejected for any variable (Kolmogorov-Smirnov, P>0.05). Discriminant analysis data were not graphed, as only a single canonical variable was obtained with two input variables. This model gave a failure rate of 2.8 % by incorrectly assigning a few of the fastest events from NGF INs as originating from FS BCs and 0 % FS BCs events as NGF IN events (overall failure rate 1.4 %). Testing this model on the total population of 320 non-contaminated events sampled from each of the 30 pyramidal neurons (9600 events) in baseline conditions, assigned 94.1 % of the events to the distribution of IPSCs originating from the FS BCs and 5.9 % as originating from NGF INs. Although this estimate is crude because all other origins are ignored, it supports the notion that spontaneous events with slow kinetics reminiscent of a NGF IN origin are rare and only contribute very little to the averaged spontaneous event.

Taken together with the insensitivity of the spontaneous event to inhibition of GAT-1 led us to speculate on the source of GABA when (assumed) synaptic-spillover was induced by electrical stimulation of afferents via a stimulation electrode. We therefore tested if the decay of the electrically evoked IPSC, induced by stimulation of all nearby afferents and sensitive to GAT-1 inhibition, also was sensitive to N-type Ca-channel blockage. The electrode was positioned at a distance of 40-50 μm from the soma along the apical dendrite of the pyramidal neuron to ensure activation of non-somatic terminals. Statistical assessment of drug effects was made on the mean (SD) of stimulation no 8-10 (baseline, BL), no 18-20 (NNC 711, 711 in short) and no 33-35 (711+ ω-conotoxin GVIA, 711+GVIA in short) using a one-way repeated ANOVA/Bonferroni (n=6). The difference in RT_10-90_ was n.s. in the three periods (1.4±0.1, 1.4±0.2 and 1.3±0.2 ms in BL, 711 and 711+GVIA, respectively, F=0.55, P=0.61). The peak was 438±207 pA in BL, 463±200 pA in 711 and 241±167 pA in 711+GVIA (F=5.3 P=0.07, Bonferroni gave P=0.03 for 711 against 711+GVIA and P=0.05 for BL against 711+GVIA). The IPSC decay, measured as τ_W_ was 12.5±5.7 ms in BL, 22.8±9.9 ms in 711 and 13.1±6.0 ms in 711+ GVIA (F=15.2, P=0.01, Bonferroni gave P < 0.004 for NNC 711 versus BL and 711+GVIA, and P > 0.76 for BL versus 711+GVIA, figure 4 F). These data indicate that electrical stimulation induce GABA release from terminals that are sensitive to both GAT-1 inhibition and N-type Ca-channel blockage. However, it seems likely that an additional source must be present for the peak to be significantly affected in conotoxin GVIA at a timecourse which is possibly too fast for NGF IN.

## Discussion

We have described some of the components involved in the increased decaytime of evoked vs spontaneous IPSCs in layer II-IV pyramidal neurons after inhibition of GAT-1 in the prefrontal cortex of adult male rats. The GABAergic output of dendritic targeting NGF INs onto pyramidal neurons is sensitive to both inhibition of GAT-1 and N-type Ca-channel inhibition by ω-conotoxin GVIA. The IPSCs originating from the FS BCs, however, were insensitive to both these treatments. We used these tools to show that electric stimulation of interneuron afferents to induce the phenomenon known as synaptic spillover in the presence of a GAT-1 inhibitor, is completely reversed by ω-conotoxin GVIA. Our findings add precision in discerning the effects of GAT-1 inhibition on GABAergic network function.

### Electrophysiological characterization of interneurons and effect of GAT-1 inhibition

The cortical NGF INs are derived from the caudal ganglionic eminence of the ventral telencephalon (Butt SJ et al. 2005; Miyoshi G *et al.* 2010) and belong to a subgroup of 5HT_3A_R-positive and VIP- and SST-negative interneurons (Olah S *et al.* 2007; Lee S et al. 2010; Rudy B *et al.* 2011). Other markers that positively label cortical NGF INs are NPY-reelin- and nNOS, but there are few exceptions to this (Jiang X et al. 2013), since it also is not clear if the hippocampal NPY- and nNOS-positive and Reelin-negative Ivy IN (Fuentealba P *et al.* 2008) exists in the cortex ((Tricoire L et al. 2010), reviewed by (Overstreet-Wadiche L and CJ McBain 2015)). The description we have given of NGF INs here, are of a late-spiking (LS) interneuron with a low input resistance, low membrane time constant which initially produce non-adapting low-frequency trains of broad FWHM APs that produce slow-rising and -decaying IPSCs which are prolonged by GAT-1 inhibition and desensitize quickly with use. These are all very-well described characteristics of NGF INs in cortical regions (Tamas G *et al.* 2003; Olah S *et al.* 2007; Povysheva NV et al. 2007; Szabadics J *et al.* 2007; Olah S *et al.* 2009; Jiang X *et al.* 2013; De Marco Garcia NV et al. 2015; Emmenegger V et al. 2018). Some features described here that appear to differ to NGF INs in other cortical regions or age of the animal, are a rather negative resting membrane potential (~ -76 mV) and a reluctance to go higher than 35 Hz in AP frequency. Also, the NGF IN phenotype in our sample of adult rat layer II-IV PFC neurons, did not appear to establish autapses, and was overwhelmingly of the LS1-type described by (Miyoshi G *et al.* 2010) in P14-21 mice, a feature that is also phylogenetically specific (Povysheva NV *et al.* 2007). Despite a similar distance between the soma of synaptically connected pyramidal neurons and NGF INs or FS BCs, typically below 20 μm, the NGF IN input to the pyramidal neuron is unlikely to overlap anatomically, because of the significant difference in the reversal potential (17 mV) for the GABA_A,SLOW_ (NGF IN) and GABA_A,FAST_ (FS BC), most likely owing to inadequate space-clamp properties of the postsynaptic pyramidal neuron (Spruston N et al. 1993; Johnston DM-SW, S. 1995; Williams SR and SJ Mitchell 2008). A distal and non-somatic input of NGF IN terminals onto cortical pyramidal neurons was also demonstrated by (Szabadics J *et al.* 2007). A central point in the functionality of the NGF INs participating in physiological network function, is the apparent discrepancy between NGF IN possible firing rate and the rapid decline of postsynaptic response, which is long-lasting (> 6 minutes) and independent of GABA_B_ and mGluR2/3 transmission. The strong diminishment of the postsynaptic response of IPSCs originating from NGF INs with increased no of APs (figure 1E-H), is not compatible with a postsynaptic origin, because we did not see such plasticity after bath application of THIP (2 μM). We assume that release from NGF IN terminals was diminished with use due to a presynaptic effect that involved inactivation of voltage gated N-type Ca-channels (below). The kinetics of the IPSCs originating from NGF INs is predominately determined by the unusually large distance of the NGF IN presynaptic terminals to postsynaptic targets, giving rise to GABA mediated volume transmission as opposed to point-to-point transmission by the FS BCs (Olah S *et al.* 2009). It is possible that dendritic filtering of distal input contribute to the slow rise time in particular, however, (Szabadics J *et al.* 2007) described that Martinotti INs produced IPSCs in pyramidal neurons with faster kinetics than NGF INs, despite a more distal dendritic input. The averaged spontaneous event resemble the events evoked from FS BC much more than the events evoked from NGF INs (figure 3 A, B), suggesting that spontaneous release from NGF INs are poorly represented in the population of spontaneous IPSCs and supported by the fact that we registered only borderline significant effect of NNC 711 on the decay of the spontaneous event (see also (Keros S and JJ Hablitz 2005)). We also argue that the kinetics of a hypothetical spontaneous event from NGF INs resemble the full evoked event, because of the very similar kinetics of the full and the scaled event in ω-conotoxin GVIA (figure 4 C, D). This lead to the estimate that the proportion of spontaneous events with rise- and decay-times that statistically assign to a possible origin from NGF IN is ~6 %. This is a crude and upper limit, as we cannot determine if this population originate from other INs very distal and low-pass filtered mono-synaptic events. However, the data support the notion that spontaneous release from NGF INs must have been rare and only contributed very little to the average spontaneous event, which also explain the very limited effect of GAT-1 inhibition on the spontaneous IPSCs. The effects of NNC 711 in differentiating between somatic and dendritic IPSCs are clear, but puzzling, because GAT-1 is highly expressed in the terminals of PV-containing FS BCs (Fish KN et al. 2011; Tricoire L et al. 2011; Armstrong C and I Soltesz 2012; Melone M et al. 2014). Instead, this separation in the effect of NNC 711 on the origin of IPSCs indicate an absence of functional extrasynaptic GABA_A_ receptors at the soma.

### N-type Ca-channels in NGF IN terminals

The demonstration that release from NGF IN terminals is sensitive to the specific N-type Ca-channel blocker ω-conotoxin GVIA is new and this toxin may serve as a tool to further characterize the functional properties of NGF INs. The N- and P/Q-type Ca-channels are both inhibited by GPCR activated Gβγ unit, which physically associate with the Ca-channel to stabilize a voltage-dependent inhibition (Herlitze S et al. 1996), but this inhibition is considerably more effective for N- than P/Q-type Ca-channels (Arnot MI et al. 2000). The pronounced use-dependent desensitization of the NGF IN-pyramidal neuron synapse which we argue is presynaptic in origin (above), but not the FS BC-pyramidal neuron synapse (figure 1), could be related to expression of specific presynaptic GPCRs, for example the NPY Y2 receptor, activated by NPY release from the NGF IN (Qian J et al. 1997; Silva AP et al. 2003; Stanic D et al. 2006) to mediate an inactivation of presynaptic N-type Ca-channels.

The event evoked by an extracellular stimulation electrode, is a concerted release of “all” terminals in a local area, and such events are typically characterized by larger decaytime than mono-synaptic spontaneous events. This is because additional GABA release cause activation of extrasynaptic receptors, a phenomenon described as synaptic overspill which is sensitive to GAT-1 inhibition (Overstreet LS and GL Westbrook 2003). The selective effect of ω-conotoxin GVIA on NGF IN release demonstrate that GAT-1 mediated increase of electrode-evoked IPSC decay (figure 4 E, F) originate from GABA released from sites that express N-type Ca-channels (NGF IN and potentially others). Since the spontaneous events were near in-sensitive to NNC 711, and since only few spontaneous events have kinetics concurrent with an NGF IN origin (~6%), taken together, these data suggest that NNC 711 and ω-conotoxin GVIA sensitive evoked synaptic overspill, is GABA release from spontaneously silent NGF INs. While the terminals of perisomatic basket cells (both PV- and CCK-expressing) is most likely activated by the stimulation electrode because of the fast IPSC rise time (~1.4 ms), the increase in the IPSC decay by GAT-1 (figure 4 E, F) is most likely dendritic in origin, because of the complete lack of response to NNC 711 at the FS BC-pyramidal neuron synapse. For this reason, it also appears unlikely that the somatic terminating CCK-containing basket cells, also expressing N-type Ca-channels (Hefft S and P Jonas 2005; Freund TF and I Katona 2007), but very little GAT-1, are involved in the increased IPSC decay induced by NNC 711. However, it is very likely that the CCK-terminals are involved in the reduction of the peak response of the evoked event after administration of ω-conotoxin GVIA. However, we cannot exclude the additional involvement of release from other dendritic targeting INs as, at this time, it is unknown which other IN terminals express N-type Ca-channels. In summary, our findings suggest that the type of synaptic overspill in the PFC that is characterized by activation of extrasynaptic GABA_A_ receptors, is synonymous with activation of spontaneously silent and dendritic terminating N-type Ca-channel expressing axon terminals, such as NGF IN terminals. Diminished expression and function of GAT-1 is a well-documented characteristic in animal models of impaired cognition as well as schizophrenic patients. Our data indicate that NGF-IN could be functionally involved in the expression of this phenotype.

## Acknowledgement

The authors declare no conflict of interest.

